# One viral sequence for each host? – The neglected within-host diversity as the main stage of SARS-CoV-2 evolution

**DOI:** 10.1101/2021.06.21.449205

**Authors:** Yongsen Ruan, Mei Hou, Jiarui Li, Yangzi Song, Hurng-YI Wang, Xionglei He, Hui Zeng, Jian Lu, Haijun Wen, Chen Chen, Chung-I Wu

## Abstract

The standard practice of presenting one viral sequence for each infected individual implicitly assumes low within-host genetic diversity. It places the emphasis on the viral evolution between, rather than within, hosts. To determine this diversity, we collect SARS-CoV-2 samples from the same patient multiple times. Our own data in conjunction with previous reports show that two viral samples collected from the same individual are often very different due to the substantial within-host diversity. Each sample captures only a small part of the total diversity that is transiently and locally released from infected cells. Hence, the global SARS-CoV-2 population is a meta-population consisting of the viruses in all the infected hosts, each of which harboring a genetically diverse sub-population. Advantageous mutations must be present first as the within-host diversity before they are revealed as between-host polymorphism. The early detection of such diversity in multiple hosts could be an alarm for potentially dangerous mutations. In conclusion, the main forces of viral evolution, i.e., mutation, drift, recombination and selection, all operate within hosts and should be studied accordingly. Several significant implications are discussed.

## Introduction

In the current practice of studying the evolution of viruses, in particular, SARS-CoV-2, each individual host is assumed to harbor one strain, or one highly prevalent strain (Candido *et al*. 2020; Forster *et al*. 2020; Rambaut *et al*. 2020; Tang *et al*. 2020). Hence, only one genomic sequence needs to be presented for all the virions within the host. In this mainstream view, the amount of diversity within each individual is negligible. Consequently, the selective advantage driving adaptive evolution would not be within the host. Instead, viral evolution would happen mainly between human hosts and the evolutionary dynamics of the virus would parallel the evolution of its human host. Viral selective advantages may take the form of surviving in the aerosol or altering host behaviors to facilitate transmission.

Despite the mainstream view, the viral evolution might happen mainly within each individual host. After all, any viral mutation should start its journey from a single virion within its host and become sufficiently common to be transmitted. Virions that proliferate more efficiently within the host would then be transmitted at a higher rate than others. For example, the spike D614G mutation can lead to more efficient replication, infection, and competition, compared with the wild-type virus (Hou *et al*. 2020; Korber *et al*. 2020). Given the large number of virions within a host, generally > 10^9^, high genetic diversity seems plausible.

If the evolution first happens within individual hosts and then continues between individuals, the two-stage evolution would add a layer of complexity and present a set of new challenges in experimentation, data collection, modeling and conceptual development. In particular, the transmission of the within-host diversity from one host to another would be a key factor. The transmission is determined by i) the number of virions transmitted in an infection (*N*_0_); ii) the number of infections by each host, corresponding roughly to R0 in epidemiology.

In the literature, *N*_0_ has often been estimated to be < 10 (and close to 1) for various viruses including SARS-CoV-2 (Popa *et al*. 2020; Braun *et al*. 2021; Lythgoe *et al*. 2021; Martin and Koelle 2021; Wang *et al*. 2021), influenza virus (Xue and Bloom 2019) and HIVs (Keele *et al*. 2008). This conclusion is usually reached by comparing the within-host diversity profiles between the donor and recipient. Such comparisons will be shown to be inappropriate for estimating *N*_0_ for SARS-CoV-2.

In this study, we will determine the level of viral diversity within each individual host by sampling the viral populations from the same individual at different times or from different tissues. If the within-host diversity is substantial, we would ask how the diversity is transformed into the between-host polymorphism. In particular, it may be possible to detect the emergence of advantageous mutations when they are still part of the within-host diversity *before* they spread among hosts. This time-lapse may be used as an early warning system. Knowing the within-host diversity is the first step in studying the long-term viral evolution.

## Results

In this study, the collection of viruses within an individual is referred to as a population or, when necessary, a sub-population. The entire collection of viruses from all infected individuals is referred to as the meta-population. Genetic diversity and polymorphism are used, respectively, for within- and between-host variation. Genetic diversity is transmitted during infection via *N*_0_ virions, which would proliferate to *N*_*t*_ virions at time t in the new host. We use the branching process to track the genetic drift during the rapid growth phase of the viral population (Chen *et al*. 2017; Ruan *et al*. 2021a; Ruan *et al*. 2021b).

### The analyses of samples collected from the same host

We first examine viral samples collected from the same host, either from the same tissue at different times or from different tissues at the same time. Strikingly, many variants are sample-specific and at a high frequency (referred to as SSH sites) as shown in Fig. 1.

**Fig. 1.**
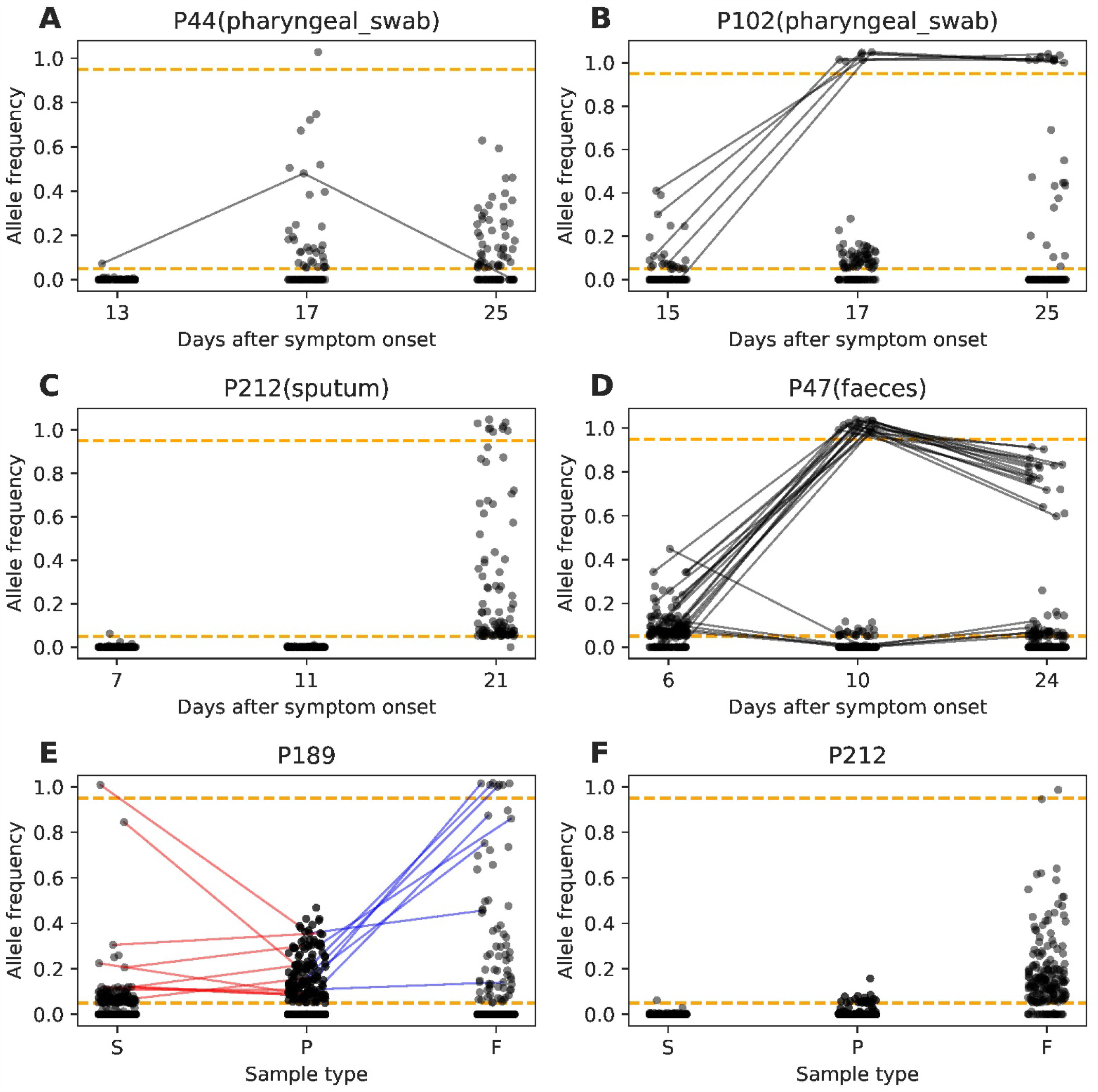
Variant frequency of samples from the same individual. Each panel represents a patient with the ID shown. The Y-axis shows the frequency of the variant (or AF, for alternative allele vis-à-vis the reference genome). The dashed orange lines show the frequency cutoff at 0.05 or 0.95. (A-D) Allele frequency from samples of the same point collected at different time points. Variants found in more than one sample are connected by a line. (E-F) Allele frequency of samples collected at the same time but from different tissues of the same patient. X-axis labels different tissue types, S for sputum, P for pharyngeal swap, F for feces. All panels show that most variants, even high-frequency ones, are sample-specific.

Fig. 1A-D compare the variants in samples collected from the same tissue of the same patient at different time points. In these panels, variants that are found in more than one sample are connected by line segments and SSH variants are unconnected by lines. Fig. 1A and Fig. 1C (from a pharyngeal and sputum sample, respectively) show a simple pattern whereby almost all variants are SSH variants with no sharing among samples. Some of these samples were collected only 2-4 days apart. Fig. 1B and 1D show a degree of sharing in addition to the SSH variants. The shared variants, however, are often very different in frequency. For example, variants that are 0.1 – 0.4 in frequency in the first sample often jump to near fixation in a second sample collected two days later and stay in high frequency in the third sample (Fig. 1B, pharyngeal). Fig. 1D (feces sample) shows a similar pattern as Fig. 1B. Such large changes are unlikely to be attributable to natural selection as they involve too many variants that change too rapidly. Furthermore, Fig. 1E-F presents the results of samples collected from the same patient at the same time but from different tissues. The differences among such samples resemble the patterns found between sample collected at different times. These differences between samples of the same individual would cast doubt on the value of comparing samples from two individuals.

### Viral sampling in relation to within-host evolution

The different patterns among samples from the same patient (Fig. 1) may suggest the within-host diversity to be far larger than realized. What would then be the total viral diversity within the same patient? And how is that diversity distributed in space and time within the host? The model presented in Fig. 2 attempts to address the issue. Briefly, each sample collected represents a small slice of the total diversity, consisting of virions recently released by a small subset of cells locally. The main viral population may be actively proliferating inside the cells unavailable to sampling.

**Fig. 2.**
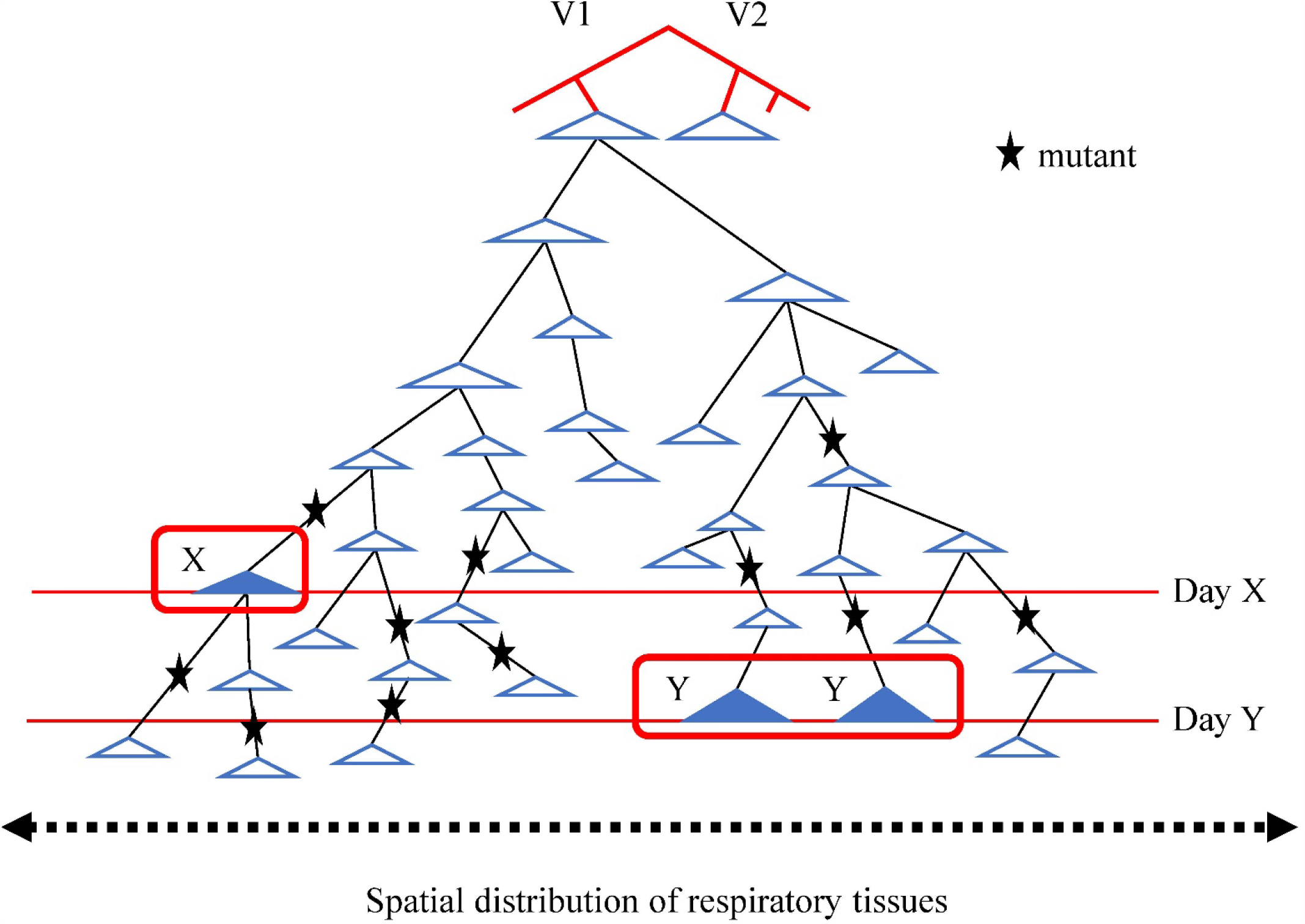
Sample collection in relation to viral proliferation within the host. The model attempts to explain the different diversity profiles between samples of the same patient. In this model, any viral sample (e.g., taken by the nasal-pharyngeal swab) represents only a small part of the geological tree that descends from a virion in the *N*_0_ cohort. The star symbol represents mutations. These virions are likely extra-cellular particles released from infected cells but have not infected another cell. Each blue triangle depicts the expansion of an ancestral virion in a local patch of tissue. It is assumed that the released virions tend to infect nearby cells. Two samples taken from the same tissue at different times (day X vs. day Y) may come from different parts of the genealogy and display very different mutation profiles. Note that the genealogy represents the descendants of only one virion. The total genealogy should consist of all lineages descending from the original cohort of *N*_0_ virions.

In this model, the viral population within a host is started by the *N*_0_ founders acquired in an infection. Each virion in the *N*_0_ sample subsequently expands into a large genealogy with numerous virions. The genealogy itself is quite different from the conventional bifurcating tree in organismal evolution. Here, the viral evolution within the host is modeled by the branching process, rather than by the conventional Wright-Fisher (WF) model (Chen *et al*. 2017; Ruan *et al*. 2021a; Ruan *et al*. 2021b).

The salient feature of Fig. 2 is that each sample represents only a small part of the genealogy, due to the transient releases of virions from the local patches of infected cells. This view of restricted sampling in both space and time may be the main reason for the discordant sample profiles from the same individual. Random sampling from the entire viral population within an individual appears untenable for SARS-CoV-2. (It may be more achievable for other viruses like HIV or HBV that has a high concentration in the peripheral blood than for coronaviruses (Koziel and Peters 2007).) The observations and modeling suggest very high viral diversity within individual hosts, within which the virus evolves.

From the genealogical tree emerge de novo mutations, some of which become SSH variants shown by the star symbol in Fig. 2. The de novo mutations, up to 50% of which could be the products of RNA editing, have been suggested to be an anti-viral strategy of the host cells. The signature of these de novo mutations is in accord with that reported for RNA editing (Di Giorgio *et al*. 2020; Mourier *et al*. 2021) with a preponderance of T-to-C, A-to-G and C-to-T changes (Fig. 3A). It can be further shown that the frequency distributions are the same for all classes of mutations (Fig. 3B), suggesting that these 12 types are comparable in their fitness consequences.

**Fig. 3.**
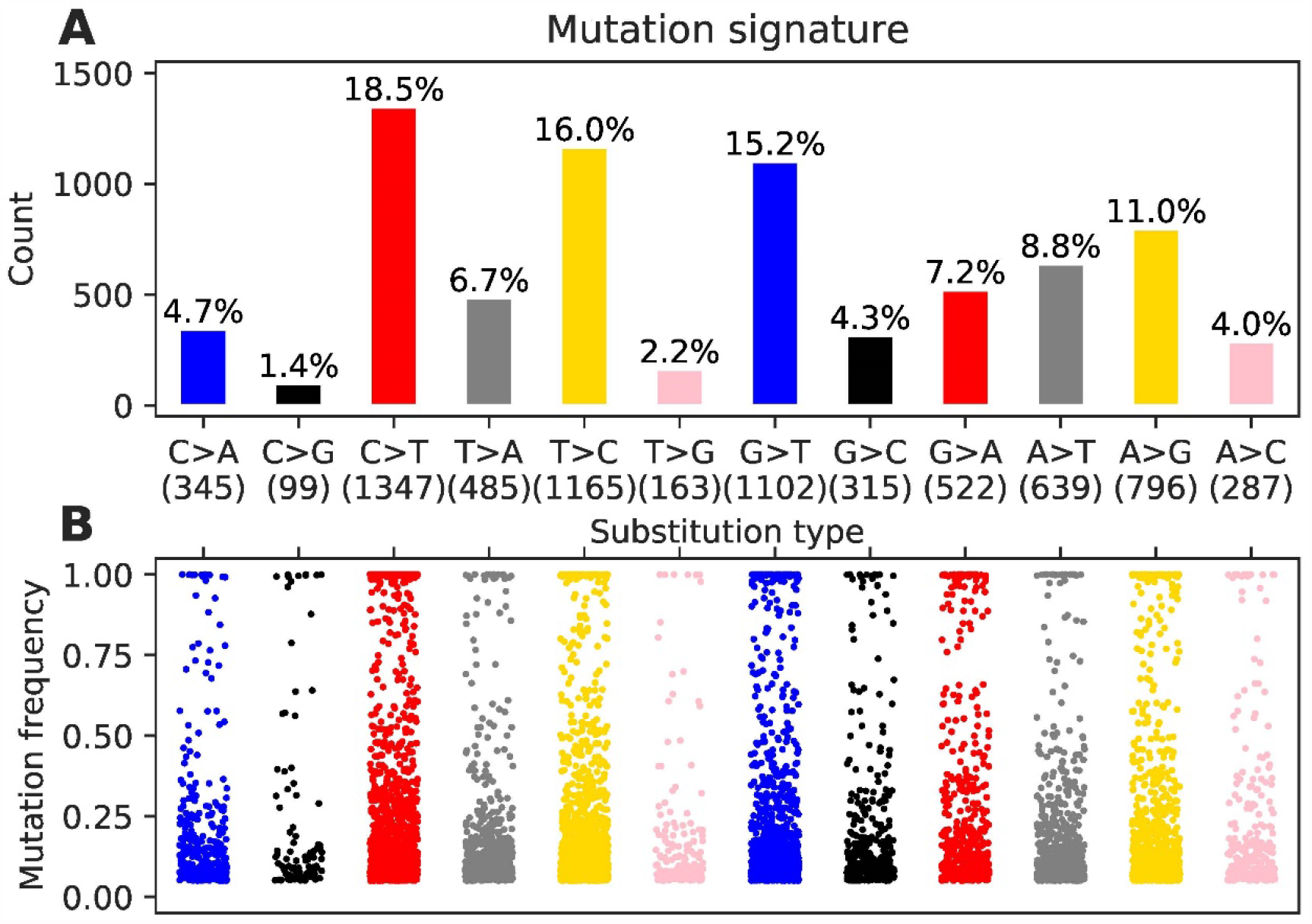
De novo mutations bearing the signature of RNA editing. (A) The distribution of the 12 types of nucleotide changes among the viral mutations, most being de novo. Each color represents the two symmetric changes, such as yellow for T-to-C and A-to-G. The yellow bars are presumably the action of ADARs (adenosine deaminases acting on RNA) while the first red bar (C-to-T) is the action of APOBEC. (B) The frequency of each mutation type in the viral samples. The frequencies appear similar across the types, suggesting comparable fitness.

### The fate of mutations within hosts

If the virus evolves within each host, what would be the fate of new mutations between their emergence and the time of transmission. Due to the rapid population expansion from a small *N*_0_ cohort, each neutral mutation would have a very low probability of being transmitted. Imagine a mutation with a fitness advantage of *s* within the host. For the neutral mutation, *s* = 0. Each of the *N*_0_ virions would grow to the size of *N*_*T*_ ≥ 10^8^. If the mutation occurs at time *t*, its frequency would be 1/*N*_*t*_ at emergence and stay near 1/ *N*_*t*_ as the population expands. A mutation is far more likely to be sufficiently frequent to be transmitted if it has a selective advantage. Fitness advantage may be of two kinds – between hosts during transmission (the B-model) or within hosts after transmission (the W-model). An advantageous mutation in the B-model would be assumed neutral within hosts. Hence, the neutral dynamics presented below applies to the advantageous mutations of the B-model.

At transmission, the number of virions transmitted to the next host is *N*_0_. Since *N*_0_ has never been estimated to be > 10^3^, the mutant frequency has to be in the order of 10^−3^ or higher to have a chance of being transmitted. The dynamics is shown in Fig. 4A as a function of the time of emergence. Fig. 4B shows that if the mutation rate is implausibly high for the first mutant to emerge when *N*_*t*_ < 100, transmission may be possible. In the W-model with *s* > 0, there is a chance for transmission even if the mutation emerges late, as long as *s* is sufficiently large (Fig. 4C).

**Fig. 4.**
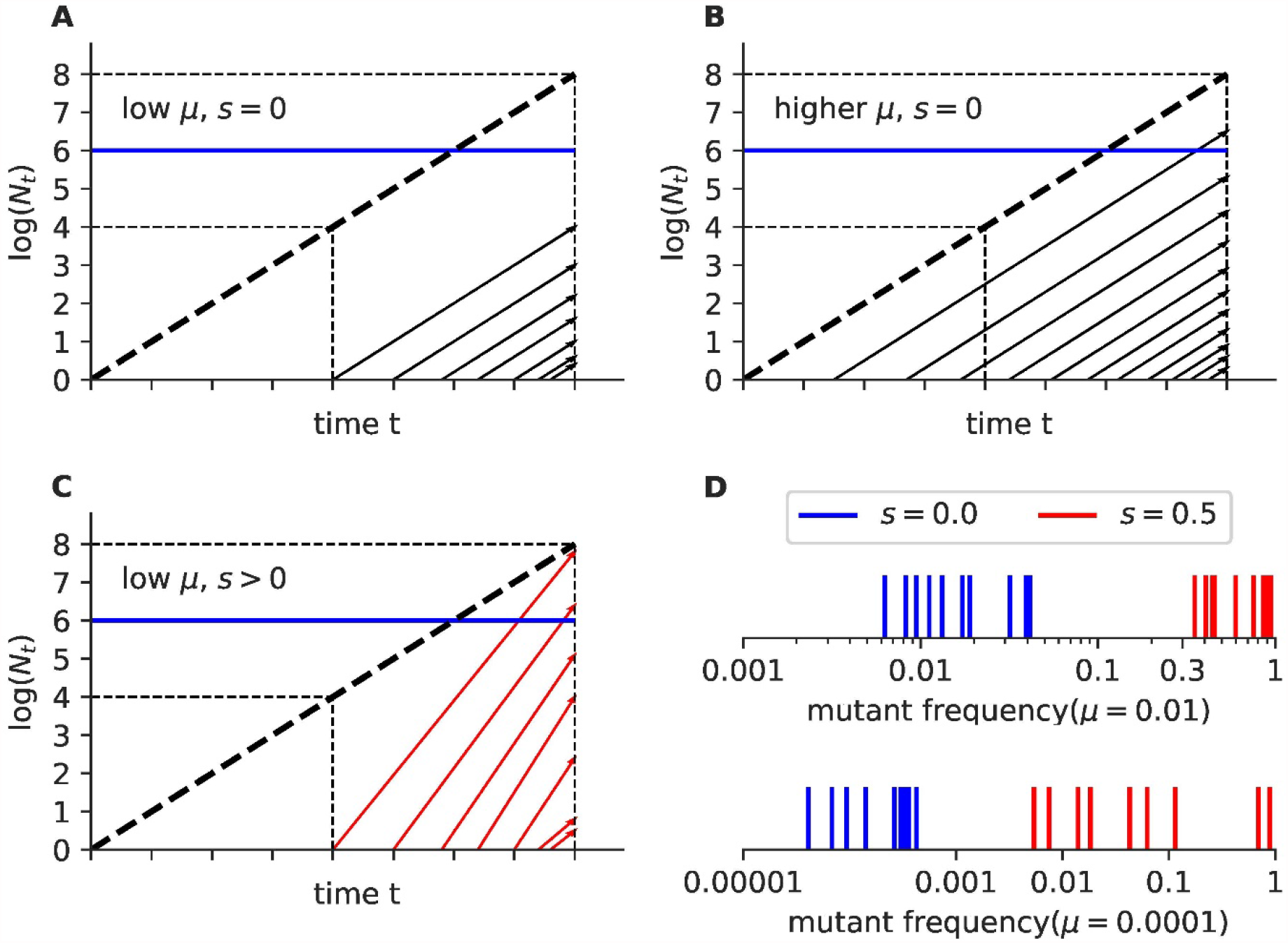
Increases of new mutations within hosts. (A-C) The increase of new mutations under different mutation rate and selection strength. Each line represents the number of virions carrying a new mutant, which increases linearly in the log scale. The lines become more closely packed in time as the population size increases and, hence, more new mutations. The blue line is the cutoff (AF = 0.01) above which a new mutation can be detected or transmitted. Note that most mutations do not rise above the blue line. (D) The mutation frequency is simulated under two different mutation rates and two selective intensities by the branching process (see Methods). The viral population grows from 1 to 10^8^. The mutation with the highest frequency at the end of the simulation in each of the 10 simulations is marked on the horizontal line. See the main text for interpretation.

Fig. 4A-4C are qualitative conjecture. We now use the branching process to obtain the quantitative results (see Methods). In a time interval (i.e. one generation), the number of virions produced by each virion is *k* with the mean of *E*(*k*) and variance of *V*(*k*). Let the mutation rate be *μ* and the mutant would reach a frequency of *x* when *N*_*T*_ = 10^8^. We let *E*(*k*) = 2 for neutral mutations and *E*(*k*) = 3 for *s* = 0.5 in the W-model. *V*(*k*) is usually set at 10*E*(*k*) as done before (Ruan *et al*. 2021a; Ruan *et al*. 2021b).The adaptive mutation rate is set at either *μ* = 0.01 or *μ* = 0.0001 per generation. These rates, artificially high for adaptive mutations, permit us to compare the fates of neutral vs. adaptive mutations.

While the mean *x* can be obtained analytically, the interest is in the outliers for the occasional mutant reaching a high *x* (say, > 0.1). The quantitative results are shown in Fig. 4D. For neutral mutations, *x* is always < 0.05 even when *μ* is extremely high at 0.01. If *μ* is set at a lower and more plausible 0.0001, *x* is always < 0.001 and *N*_0_ has to be > 1000 for transmission. For *s* = 0.5 with *μ* = 0.01, *x* is always > 0.3 and often > 0.5; hence, *N*_0_ >= 3 should be sufficient for the mutation to be transmitted. When *μ* = 0.0001, there is a 20% chance that *x* > 0.5 and the mutation is still highly transmissible.

We should note that the parameters of *μ* = 0.0001 and a selective advantage of *s* = 0.5 (i.e., a strongly advantageous mutation emerges once among 10,000 virions) are likely too high as well. The high values are used to show how adaptive mutations within hosts would greatly speed up the rate of viral evolution. At a lower mutation rate, there would be almost no chance for neutral mutations to reach a high frequency to be transmitted. For the W-model, the rate of adaptive evolution may still be small but it would be non-zero if s is not too small.

### From within-host diversity (iSNVs) to between-host polymorphism (SNPs)

The evolution of virus in the meta-population happens in stages as illustrated in Fig. 5A where each circle portrays the viral diversity of an infected individual. Stage 1a and 1b are mainly about within-host evolution while Stage 2a and 2b are about between-host evolution. Multiple circles in a stage comprise the meta-population. The evolution starts with iSNVs (for intra-host nucleotide variation), which may or may not evolve into SNPs (single nucleotide polymorphism in the host population). All SNPs obviously evolve from some iSNVs but most iSNVs probably fail to become SNPs. In the current practice, only the prevalent variant is presented for each infected individual, shown next to the circle.

**Fig. 5.**
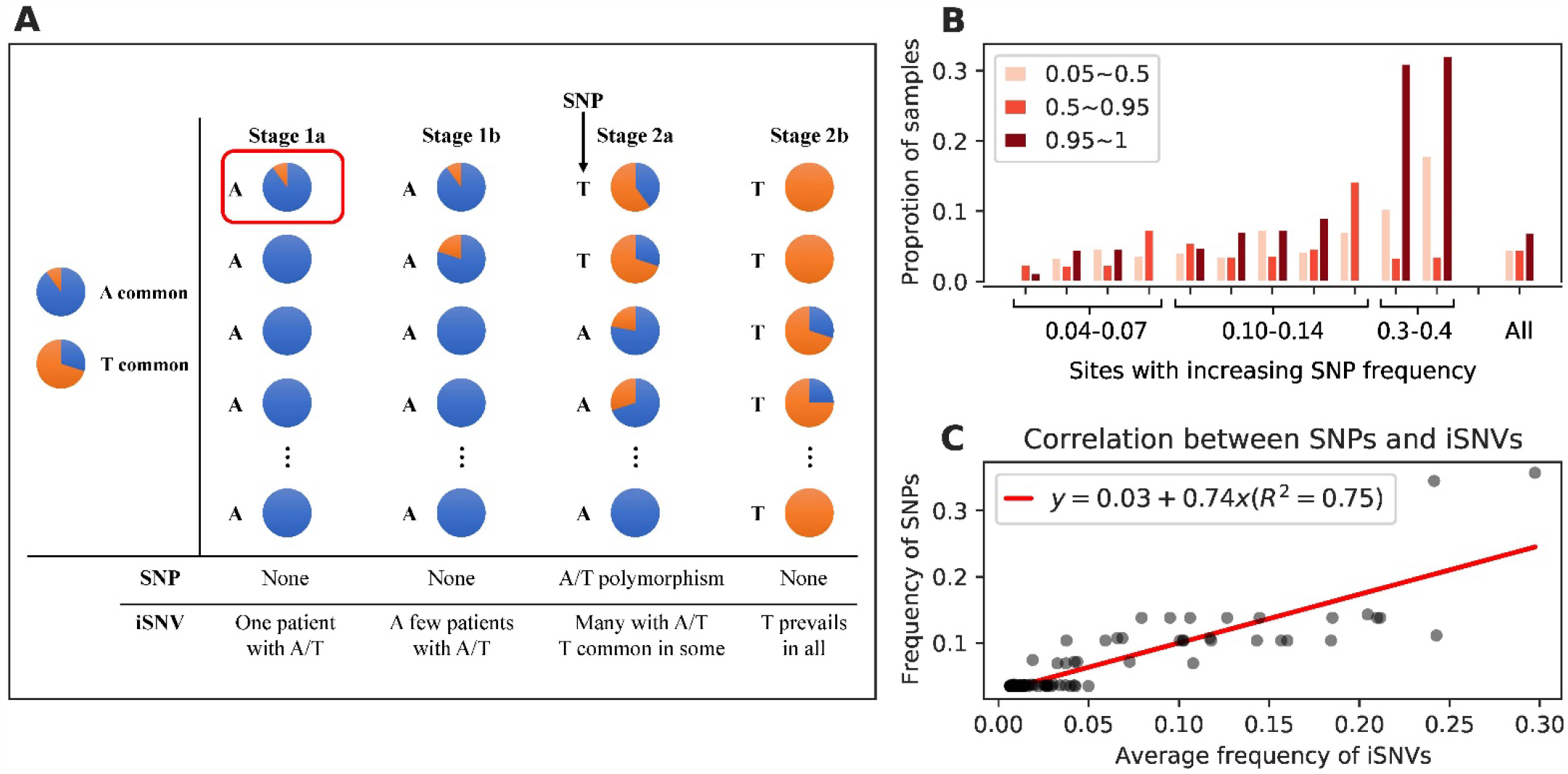
Relationships between within-host diversity and between-host polymorphism. (A) A model connecting the within- and between-host evolution in 4 stages (1a, 1b within hosts and 2a, 2b between hosts). Each within-host population is represented by the prevalent variant in the analysis of SNPs. State 2a is the only stage where iSNV and SNP are both observed. (B) The proportion of samples whereby the variant allele is lost, fixed or in the frequency range of either (0.05 – 0.5) or (0.5 – 0.95). (C) The correlation between the SNP frequency on the Y axis and the mean iSNV frequency on the X-axis. The two quantities are correlated, as expected.

In stage 1a, a T-mutant emerges from the viral population of A in an infected individual (the red-border box). Stage 1 is likely the most common outcome because the new mutant is unlikely to rise to a high frequency to be transmitted. Occasionally, if T has a selective advantage, it will rise to a “transmissible” frequency, which depends on the size of *N*_0_ (see below) and the evolution progresses to stage 1b. Between stages 1a and 1b is the most crucial transition when the mutant comes “out of the gate”. From stage 1b and on, T would increase in frequency is some individuals while the A/T diversity would continue to infect more individuals as illustrated in Fig. 3. Stage 2a is the only stage when SNPs overlap with iSNVs. Stage 2b is the next stage of evolution but appears to be the mirror image of Stage 1b, neither of which has the SNP polymorphism.

The stage of greatest interest is Stage 2a which is reflected by the data on the patient cohort of Beijing. Fig 5B shows the proportion of samples where the iSNV frequency is < 0.05, 0.05 – 0.5, 0.5 – 0.95 and > 0.95, respectively. Each quartet of bars represents SNPs of an increasingly higher frequency as marked at the bottom. It is evident that iSNVs and SNPs move in the same direction and the number of samples with iSNVs is substantial. Across all SNP sites (the last set of bars), 8.6% of samples are in the transition from iSNV to SNP whereas fixed SNPs account for only 6.8% of the samples. The mean mutant frequency from all samples for each site, iSNV (mean), is plotted against the SNP frequency (Fig. 5C). The high correlation coefficient (R^2^ = 0.75) supports the view that SNP increases in frequency as iSNV becomes more common.

### The transmission of diversity from donors to recipients

The genetic diversity shaped by events within hosts may or may not be transmitted to other hosts, depending on how often the transmission happens and how many virions are transmitted each time. The former can be roughly estimated since it is correlated with basic reproduction number (*R*_0_). The latter, i.e., the size of *N*_0_, has been of substantial interest lately. *N*_0_ is usually done by comparing the diversity profiles between donors and recipients.

Not unexpected, two samples from the same patient shown in Fig. 6A look like samples from the donor-recipient pairs. Fig. 6B presents a typical example of the observed pattern between donors and recipients. This example is a composite of multiple samples from patients of the Beijing area in the period of January to April 2020. Another example (Fig. 6C-6D) is a re-compilation of the published data in Austria (Popa *et al*. 2020; Martin and Koelle 2021). In all these panels, variants common in one host are often undetectable in the other (i.e., data points on the X or Y axis).

**Fig. 6.**
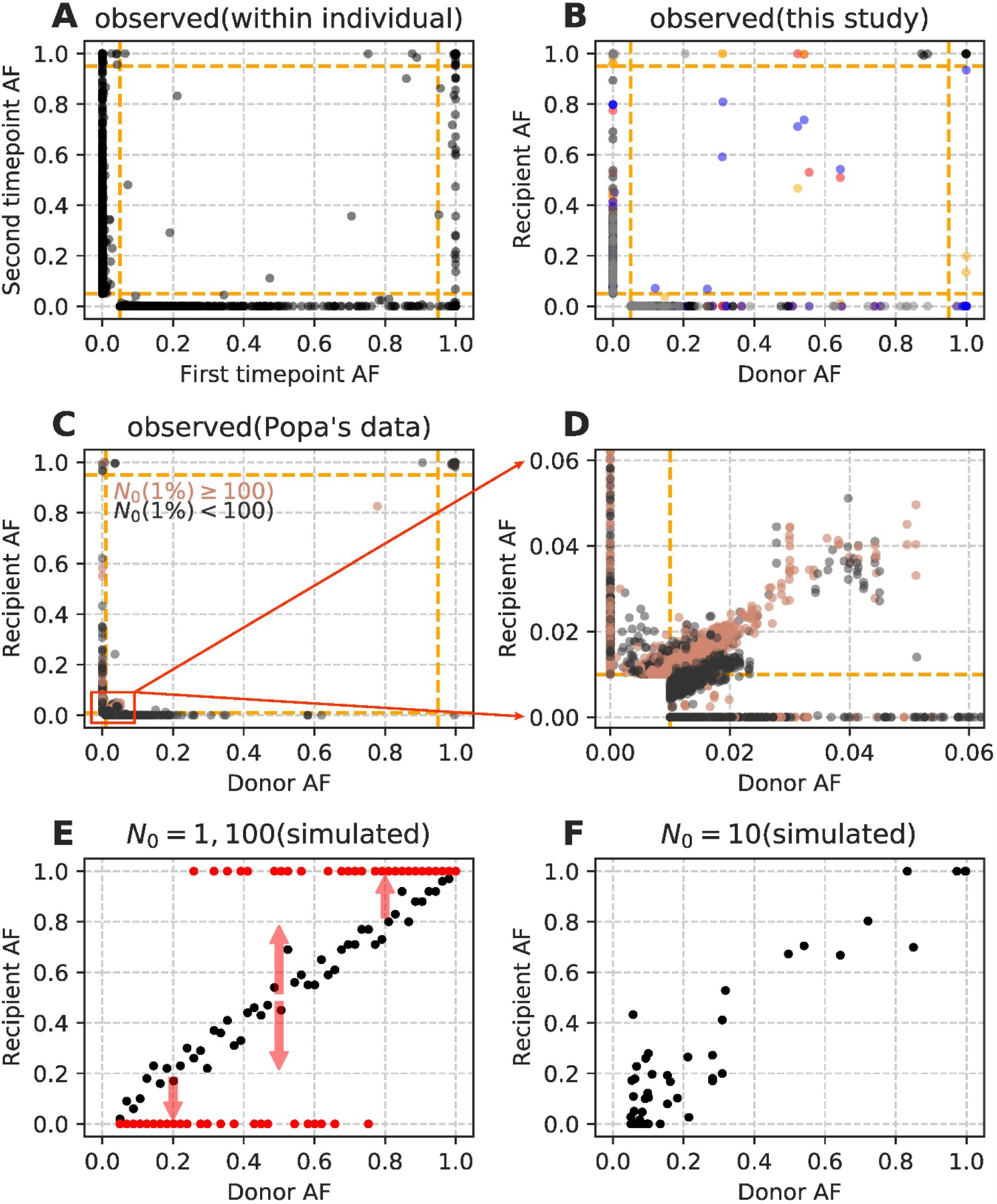
Variant frequencies between donors and recipients. The X axis shows the allele frequency (AF) in the donors and the Y axis shows that of the recipients. In all panels, data points are the composite of multiple pairs of comparisons. (A) Allele frequency comparisons between samples collected at different times from the same patient for comparisons with the other panels. (B) The composite donor-recipient comparisons from 5 pairs of family members. Each donor-recipient pair is represented by a different color. Data points falling on the X- or Y-axis are SSH (Sample Specific High-frequency) variants, which have been commonly reported but rarely analyzed (Popa *et al*. 2020; Martin and Koelle 2021; Wang *et al*. 2021). (C-D) Allele frequencies of 39 donor-recipient pairs reported in Ref. (Popa *et al*. 2020). (E) The expected donor-recipient relationship by simulations for *N*_0_ of 1 (red points) and 100 (black points). The arrows indicate the distribution of fixed or lost variants when *N*_0_ = 1. The allele frequency in the donor is taken from a uniform distribution. (B) *N*_0_ = 10. Here, the variant frequency in the donor is taken from the empirical observations of Fig. 2B of Ref. (Lythgoe *et al*. 2021). Genetic drift during the proliferation in the host is simulated by the branching process model of Ref. (Chen *et al*. 2017; Ruan *et al*. 2021a; Ruan *et al*. 2021b).

The large donor-recipient difference has led to the conclusion of a very small *N*_0_ during transmission (Poon *et al*. 2016; Mc Crone *et al*. 2018; Xue and Bloom 2019; Popa *et al*. 2020; Braun *et al*. 2021; Lythgoe *et al*. 2021; Martin and Koelle 2021; Wang *et al*. 2021). However, only a certain type of differences can be explained by a small *N*_0_. In the simulations of Fig. 6E, the donors and recipients are shown to have similar profiles when *N*_0_ = 100. With *N*_0_ = 1, the donor-recipient difference is large but the variants are either fixed or lost with the arrows indicating where the data points would be. At *N*_0_ = 10, the pattern is closer to *N*_0_ = 100 than to *N*_0_ = 1 (Fig. 6F). The difference between the expected (Fig. 6D-6F) and the observed (Fig. 6B-6D) would argue against transmission bottlenecks of any size.

It seems obvious that SSH variants have to be discounted in estimating *N*_0_. In Fig. 1E, we re-analyzed the data of Popa et al. It can be seen clearly that, by removing SSH variants entirely, the pattern agrees with Popa et al.’s estimates of *N*_0_ >100 for SARS-CoV-2 (Popa *et al*. 2020; Martin and Koelle 2021). Nevertheless, the accuracy of estimation based on the donor-recipient comparisons would be low if *N*_0_ > 10. At present, a conservative conclusion would simply be that *N*_*0*_ *>> 10*.

## Discussion

The 2-stage evolution of viruses has several implications. Let the average number of virions in a host be *N* and the number of infected individuals in the meta-population be *M*. The size of the meta-population is hence *MP* = *N* × *M*. For SARS-CoV-2, *MP* is easily 10^15^ or larger at some point.

First, given the extremely large *MP*, the rate of adaptive evolution of SARS-CoV-2 may in fact be quite modest. In a 2-stage system, a mutant has to clear the hurdle within the host in order to enter into the meta-population. Thus, moderately advantageous mutations may not be able to rise to a frequency high enough to be transmitted. On the other hand, although strongly advantageous mutations may be able to rise, they are generally uncommon. In a sense, the two stages of viral evolution divided by inter-host transmissions put a brake on the rapid evolution of viruses.

Second, the speed constraint on viral evolution could be broken when *M* becomes large. The increase in *M* can speed up the rate of adaptive evolution (*R*) with the standard population genetic expression of

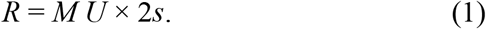

Here, *U* is the rate of adaptive mutation and *s* is the selective advantage (Eyre -Walker 2006). Eq. (1) is as conventionally understood but, in viral evolution, an increase in R would also lead to an increase in M. In other words, R and M would form a positive feedback loop as a larger R (more advantageous mutations) would lead to a larger M and vice versa. Thanks to the positive feedback look, *M* could be in a runaway process, akin to the mutation accumulation of cancer cells (Ruan *et al*. 2020). Pandemics are not as common as one may fear because the feedback loop cannot easily get started. Nevertheless, when *M* is allowed to become very large (due to host behaviors, for example), the loop may be able to get rolling thus kicking start the runaway process. Whether COVID-19 has entered the runaway process at some point of the pandemic will be a worthy topic for the future.

Third and fortunately, there may be a way to catch the rise of an advantageous mutation before it is too late. While stage 2a of Fig. 5 is a most dangerous stage, an advantageous mutation may stay in stage 1b for a while before it advances to stage 2a. When an advantageous mutation reaches stage 2a, it would have spread too far to hold back. The D614G (Hou *et al*. 2020; Korber *et al*. 2020; Plante *et al*. 2021; Volz *et al*. 2021) and N501Y mutations (Dejnirattisai *et al*. 2021; Deng *et al*. 2021) may be such examples. In the example of Fig. 5A, if we could detect T in several infected individuals as a low frequency iSNV before it progressed to stage 2a, it may be possible to stop T from getting out of the gate. It thus behooves us to have some within-host diversity data. The solution needs not be to increase the number of samples, which may be logistically difficult. Instead, all samples could be sequenced to a greater depth and in high fidelity such that all minor variants at 1%, or even lower, can be detected with certainty. The diversity data within hosts would then be immensely more informative with only modest extra efforts. Virion samples of the respiratory tract are of the greatest interest.

In conclusion, the evolution within hosts is the very first step of the long-term evolution of viruses. Figs. 4-5 show how adaptive mutations would spread in the meta-population. This spread is key to our view of the evolutionary dynamics and, hence, the containment strategy. The standard data of one viral sequence per host do not lend themselves easily to within-host analyses. Fortunately, slightly greater efforts in bulk sequencing, data presentation and model building may fill the large gap in studying SARS-CoV-2 as well as many other viruses.

## Methods

### Samples

Our study included 60 COVID-19 patients of 23 family clusters between January 20 and April 20, 2020 in Beijing Ditan Hospital. Totally there are 135 samples (47 pharyngeal swabs, 59 sputum, and 29 fecal samples), including 34 samples of 16 imported patients (oversea travelers) in 6 families and 101 samples of 44 local patients in 17 families. There are 5 donor-recipient transmission pairs, confirmed by epidemiological data (see Data file S1).

### Sequencing

Viral RNA was extracted using the QIAamp® ViralRNA Mini Kit according to the manufacturer’s instructions. After performing rRNA removal using the MGIEasy rRNA Depletion Kit (BGI, Shenzhen, China), we used the novel Metagenomic RNA (Chen *et al*. 2020). The final viral-enriched libraries were sequenced on an Illumina NextSeq500 in 2×75bp pair-end mode.

### iSNVs calling

(1) After quality control, sequencing reads were pair-ended aligned to the reference genome sequence (GenBank accession no. NC 045512.2 (Wu *et al*. 2020)) using Bowtie2 v2.1.0 (Langmead and Salzberg 2012) by default parameters and the alignments were reformatted using SAMtools v1.3.1 (Li *et al*. 2009); (2) for each site of the SARS-CoV-2 genome, the aligned low-quality bases and indels were excluded to reduce possible false positives and the site depth and strand bias were recalculated; (3) samples with more than 3,000 sites with a sequencing depth ≥100× were selected as candidate samples for iSNVs calling.

A series of criteria were used to ensure high quality iSNVs with Q20 reads support: (1) minor allele frequency of ≥ 5% (a conservative cutoff based on an error rate estimation); (2) depth of the minor allele ≥ 5. After calling variants, we used ANNOVAR software (Yang and Wang 2015) to annotate the variants and found the count of alternative allele and total depth for each variant using SAMtools (see Data file S1).

### Reanalysis of previously published SARS-CoV-2 data

We reanalyzed 138 COVID-19 samples with clinical information of Popa’s data (Popa *et al*. 2020), which including 39 transmission pairs. We downloaded the clinical information and vcf files at https://zenodo.org/record/4247401 and https://github.com/m-a-martin/sarscov2_nb_reanalysis. We used python scripts to merge the frequency of iSNVs of these 138 samples (see Data file S2). For each transmission pair, we identified the variants at frequency of ≥ 1% and showed the allele frequency change between donor and recipient (Fig. 6C, 6D). We used the threshold 100 of transmission bottleneck (*N*_0_), estimated by Martin et al (Martin and Koelle 2021), to divide the alleles into two groups (see Fig. 6C, 6D).

### Genetic drift when population grows (*E*(*k*) > 1)

Based on branching process, Chen et al. obtained the genetic drift after single generation (Chen *et al*. 2017). Here we expand it and get the genetic drift after multiple generations, which can be used to estimate the variance of alternative allele frequency within host. According to (Chen *et al*. 2017), the average and variance of population size at time *t* are

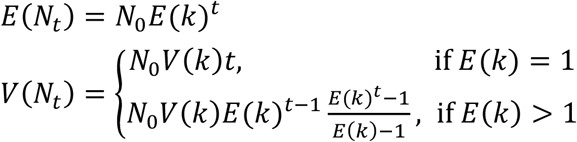

Assuming there are two kinds of alleles, and their numbers at generation *t* are *I*_*t*,_ *J*_*t*,_ *I*_*t*_ and *J*_*t*_ will be independent. If there is no selection, then

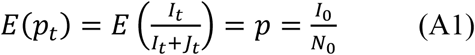

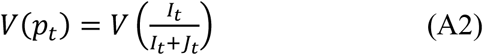

According to bivariate first order Taylor expansion (Duris *et al*. 2018), when *E(k) > 1*

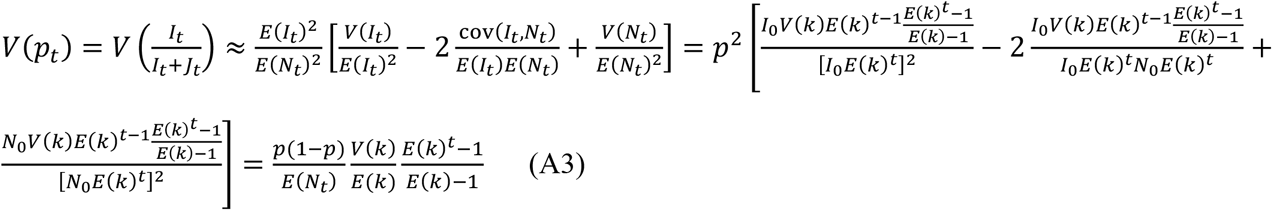

Specially, when *t=1*,

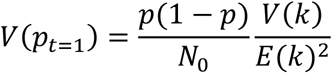

which is the same as Eq. (5) in (Chen *et al*. 2017).

### Expected allele frequency after transmission

Assuming there are *n* alleles with corresponding frequencies *x*_*1*_,*x*_*2*_, *…, x*_*n*_ in donor, we will obtain the expected allele frequency of recipient under a particular transmission bottleneck size *N*_*0*_ as follows.

For traditional WF model (Fig. 6E), each allele is independent and its allele frequency in next generation will follow binomial distribution. Thus, given transmission bottleneck size *N*_*0*,_ for the allele with frequency *x*_*i*_ in donor, its frequency in donor, *x*_*i*_*’* will be sampled from following binomial distribution

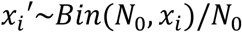

After transmission, we assume the virus will grow to a particular number, *N*_*t*,_ before it be sampled and sequenced (Fig. 6F). During the branching process of virus growth, we assume each virus will generate *k* number of offspring, where *k* follows a negative binomial distribution with mean *E*(*k*) and variance *V*(*k*). Thus, the expected time at which the virus is sampled to determine the recipient allele frequency (denoted by *x*_*t*_) is

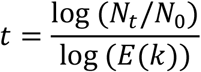

According to Eq. (A1) and Eq. (A3), given the initial allele frequency *x*_*i*_*’*, we can obtain the mean and variance of *x*_*t*_ when population size grows from *N*_*0*_ to *N*_*t*_:

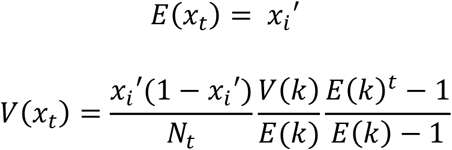

Simply, we can assume *x*_*t*_ follows a normal distribution with mean and variance to be *E(x*_*t*_) and *V(x*_*t*_) Now, we can obtain the expected allele frequency in donor recipient by sampling from the normal distribution.

### Simulating the fate of new mutations within host under different selection strength

As done before (Ruan *et al*. 2021a; Ruan *et al*. 2021b), we simulate mutation accumulation of virus within host by branching process. In the branching process, each virion would produce *k* descendants in a period of time, say 24 hours, with the mean and variance of *E*(*k*) and *V*(*k*), respectively. *E*(*k*) is the rate of viral proliferation and can be estimated with reasonable accuracy. In the WF model, *V*(*k*) would equal *E*(*k*) and is unlikely to be correct for viruses. Instead, the population genetic parameters of SARS-CoV-2 (and perhaps many other viruses as well) suggest that *V*(*k*) may be far larger than *E*(*k*). For example, it has been estimated that only 10^5^ – 10^7^ cells in a human host are infected at any moment (Sender *et al*. 2020). This is a tiny fraction of human cells, given the estimated 10^13^ – 10^14^ cells in the host. Hence, each infected cell may need to produce a large number of virions. The numbers suggest a very large *V*(*k*) for SARS-CoV-2. Here we let *V*(*k*) = 10*E*(*k*), where *k* follows negative binomial distribution. The initial *E*(*k*) is 2, which can be increased by new adaptive mutations.

First, we set up a mutation-free viral population (initial population size is 8 to avoid population extinction) at initial time. Mutations were accumulated in a Poisson process with mutation rate μ = 0.01, 0.0001 per generation until the population size reached 10^8^. In neutral evolution, we assume each new mutation is neutral (*s* = 0). And *s* is 0.5 for each mutation in adaptive evolution. For each parameter set, we repeat the simulation 10 times (see Fig. 4D). Finally, we calculated the frequency of the most dominant allele (i.e. the highest frequency of all the accumulating mutations).

### Relationships between within-host diversity and between-host polymorphism

To estimate the correlation between SNPs and iSNVs frequency, we picked 29 of 44 local patients (101 samples in total) by following criteria: (1) patients with at least one sputum or pharyngeal samples; (2) the 100X genome coverage ratio of samples in this patient is greater than 80%. For the patient with more than one samples fitting the criteria, we only picked the sample with the highest coverage. Then, we used the 29 local samples (one sample for one patient) to find SNP sites based on the frequency of iSNV sites.

For each iSNV site in the 29 samples, we counted the number of samples whose iSNV frequency is < 0.5 (reference allele), denoted by *N*_ref_. Similarly, we denoted the number of samples with iSNV frequency ≥ 0.5 by *N*_alt_ (alternative allele). If *N*_ref_ > 0 and *N*_alt_ > 0 (reference allele and alternative allele existed at the same time), the iSNV site corresponds to a consensus SNP site. And the SNP frequency was *N*_alt_ / (*N*_alt_ + *N*_ref_). Based on this, we found 95 SNP sites in the 29 samples. For the 95 SNP sites, we also calculated the average iSNV frequency in the 101 local samples (see Data file S1). The correlation between SNP frequency and iSNV frequency is shown in Fig. 5C.

